# ILORA: A database of alien vascular flora of India

**DOI:** 10.1101/2021.05.28.446252

**Authors:** Vidushi Pant, Chinmay Patwardhan, Kshitij Patil, Amiya Ranjan Bhowmick, Abhishek Mukherjee, Achyut Kumar Banerjee

## Abstract

1) Biological invasions pose an unprecedented threat to biodiversity and ecosystems at different spatial scales, especially for a biodiversity-rich developing nation like India. While country-level checklists of alien taxa are important, databases having their biological and ecological attributes are of paramount importance for facilitating research activities and developing policy interventions. Such a comprehensive database for alien flora is lacking in India.

2) We have curated data for 14 variables related to ecology, biogeography, introduction pathway, socio-economy and distribution of 1747 alien vascular plant species from 22 national and global sources to produce the Indian Alien Flora Information (ILORA) version 1.0 database. This paper describes the detailed methodology of curating these data along with the rationale behind selecting these variables.

3) The database, the first of its kind for the Indian alien flora, will provide easy access to high quality data and offer a ready reference to comprehend the existing scenario of alien plant species in the country. The database is dynamic and will be updated regularly. It has a provision to incorporate user submitted data, which will allow increasing the resolution of the database as well as the expansion of its capacity.

4) The database is envisaged to become a nationwide collaborative platform for a wide spectrum of stakeholders. It is freely accessible via an online data repository as well as through a dedicated website (https://ilora2020.wixsite.com/ilora2020).

## 1. Introduction

Biological invasion has been described as a defining feature of the Anthropocene (van Kleunen et al., 2015). The emerging economies, like India, are facing the greatest threat of invasion as a consequence of increasing globalization of trade as well as the changing climate (Lenzen et al., 2012). This necessitates the creation of a country-level repository of alien flora as the first step towards identifying emerging invaders and managing the ones that have already been established (Pagad et al., 2018). While country-level checklists of alien flora exist in India [e.g. (Khuroo et al., 2012)], the biological and ecological attributes of these alien plant species remain scattered across a multitude of databases rendering their access tedious. Some of this information, although retrievable from global databases, often lack resolution at the national or regional levels (but see The Global Naturalized Alien Flora database, hereafter GloNAF; (van Kleunen et al., 2019). Further, data extraction often involves intensive and informed query-directed search processes to garner species specific information. Easy access to such information at a finer scale is of paramount importance for developing country-level policy interventions and facilitating research and development concerning the alien species of a country. Such a comprehensive database is lacking in India (but see (Inderjit et al., 2018).

Here, we introduce the **I**ndian A**l**ien Fl**o**ra Info**r**m**a**tion (ILORA) Version 1.0 Database, a dynamic and open-source repository of comprehensive information on taxonomy, introduction pathway, use, and biogeography of 1747 alien vascular plant species of India. Our intention was not to create another country-level checklist, rather we envisage that ILORA will assist in novel research on alien plant invasions in India and can be transformed into a nationwide collaborative platform for stakeholders.

## 2. Materials and Methods

The database formation included two major steps – 1) categorization of the alien flora, and 2) data curation for 14 variables of these species. The steps have been explained in brief in the following text along with the overview of the data files (Table 1). The detailed methodology (including resources and search criteria) and the future plans have been provided as supporting information.

**Table 1.** Overview of the data records of ILORA version 1.0 representing the contents and data type for each of the twenty data files

|  | **File name (with extension)** | **Data contents** |
| --- | --- | --- |
| 1 | ILORA_1_SpCategorization.csv | The reported origin and invasion status of the alien species present in India |
| 2 | ILORA_1a_Sp.Categorization.Metadata.csv | Metadata for determining origin and invasion status |
| 3 | ILORA_2_GeneralInformation.csv | Common and vernacular names, life form, duration, and type |
| 4 | ILORA_3_NativeRange.csv | Native range at TDWG* Level 2 |
| 5 | ILORA_4_Introduction.csv | Month and year of first reported occurrence in India; introduction pathway(s) |
| 6 | ILORA_5_EconomicUses.csv | Economic uses at TDWG* Level 1 |
| 7 | ILORA_6_MarketDynamics.csv | Price and number of plants/seeds sold in two online nurseries of India |
| 8 | ILORA_7_Habitat.csv | Potential habitat(s) |
| 9 | ILORA_8_NaturalizedRange.csv | Naturalized ranges at global and Indian scale at TDWG* Level 4 |
| 10 | ILORA_9_Occurrence.csv | Latitude and longitude data |
| 11 | ILORA_10_Distribution.csv | Distribution in Indian states and union territories |
| 12 | ILORA_11_LULC.csv | Land-Use and Land-Cover categories inhabited |
| 13 | ILORA_12_Anthromes.csv | Indian anthromes inhabited |
| 14 | ILORA_13_Ecoregions.csv | Indian ecoregions inhabited |
| 15 | ILORA_14_Climate.csv | Köppen-Geiger climate classes of India inhabited; the temperature and precipitation data |
| 16 | ILORA_15_Summary.csv | Status (presence/absence) of data for each of the above mentioned 14 data files for each of the alien plant species |
| 17 | ILORA_16_Cultivated.Species.csv | Standardized species names of cultivated aliens |
| 18 | ILORA_17_R.Codes.txt | R codes used to extract variable information |
| 19 | ILORA_Methods.pdf | Document having information on rationale of variable selection, data sources, search queries; future scopes of data enrichment |
| 20 | ILORA_HowToRead.pdf | Document having information of the data types recorded in columns in each of the data file |

\* TDWG: International Working Group on Taxonomic Databases for Plant Sciences

### 2.1 Categorization of the alien flora

We created a comprehensive list of alien plant species of India from three national checklists, viz. Alien flora of India (1599 species; (Khuroo et al., 2012), Naturalized alien flora of the Indian states (499 species; (Inderjit et al., 2018), and the GRIIS- India Version 1.3 (2082 species; (Sankaran et al., 2020). The species names were standardized using the *WorldFlora* package (Kindt, 2020) in R version 4.0.2 (R Core Team, 2020). The duplicates and synonyms, infraspecific taxa and artificially hybridized species were removed.

The selected plants (n=2503) were subjected to a two-step verification process (Fig.1) to determine their – i) origin status (native or alien to India) and ii) invasion status (degree of naturalization) (Pyšek et al., 2004). Briefly, we first identified the cultivated aliens (n=756) from the national and global databases. These species were retained in the database; however, no variable information was extracted for these species. The origin status of the remaining species (n=1747) was ascertained using global databases. If the origin status of a species was identified as ‘native’ or ‘alien’ to India in at least one of these databases and was not contradicted by the others (n=1576), it was accepted. However, in case of any contradiction between databases (n=162), origin status was determined through literature search. The species with ambiguous origin status were categorized as ‘cryptogenic’ (i.e., uncertain biogeographic status) following Essl et al. (2018). Therefore, the origin status of the alien species was categorized as – alien (n=1388), native (n=334) and cryptogenic (n=25). From the list of alien species (n=1388), the ‘invasive’ species were categorized by consulting the GRIIS database (n=220). The species which have been identified as naturalized aliens by Inderjit et al. (2018) and the GloNAF database, were categorized as ‘naturalized’ (n=237). The remaining alien species were identified as ‘casual alien’ (n=931) following Khuroo et al. (2012). Finally, a confidence score was attached to each of the species’ categories. It is important to note here that given the vast geographic extent of India, it is possible that some of the species identified as ‘native’ may be native in one part of the country and introduced to others. The status of these species may change by availability of regional-scale data and therefore, we included detailed information of them in the database.

**Fig.1:**
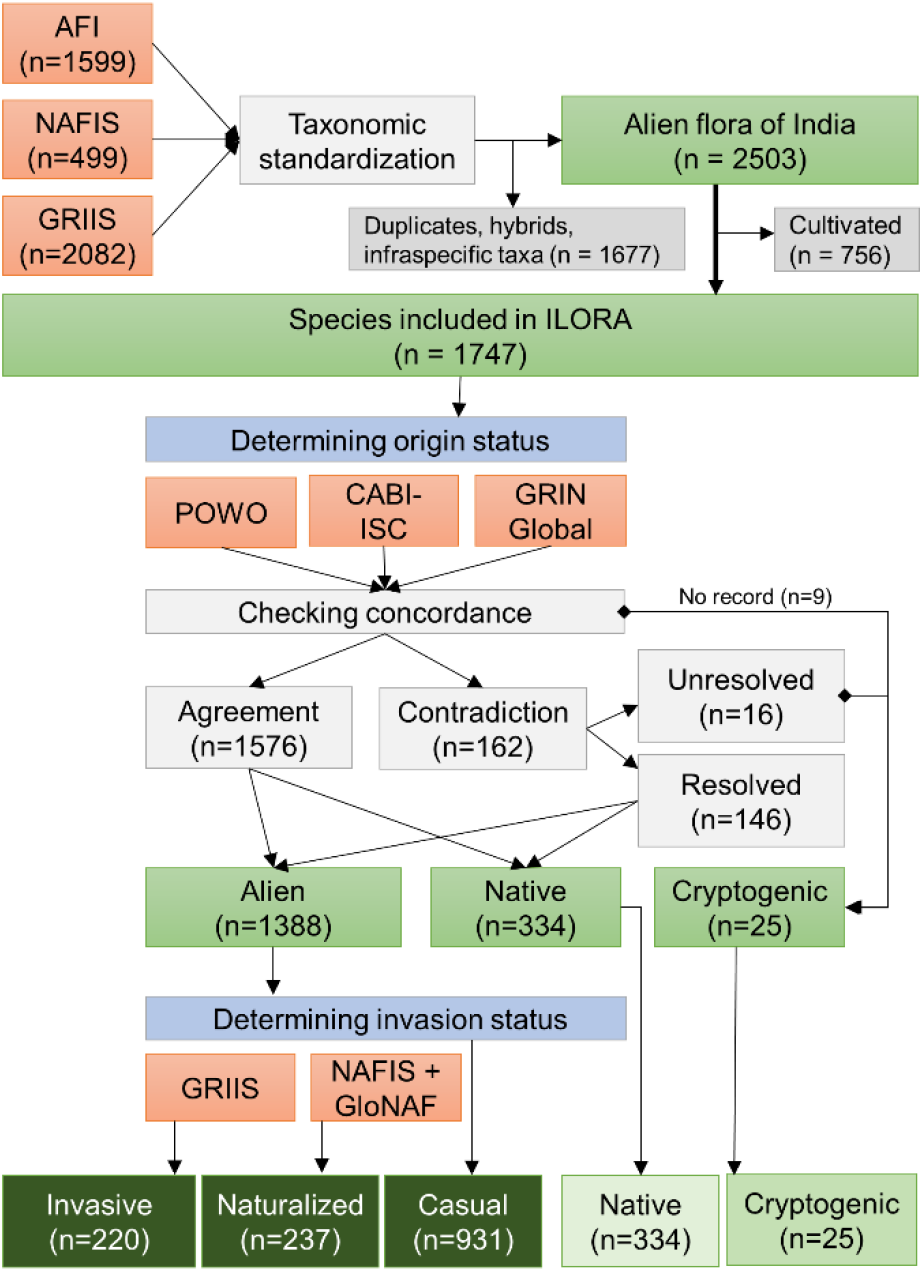
The methodology followed for the categorization of the Indian alien flora [AFI – Alien Flora of India (Khuroo et al., 2012); NAFIS: Naturalized Alien Flora of the Indian States (Inderjit et al., 2018); GLONAF: The Global Naturalized Alien Flora database (van Kleunen et al., 2019); GRIIS: Global Register of Invasive and Introduced Species (Sankaran et al., 2020)]

### 2.2 Data curation

#### 2.2.1 Taxonomy and general information

For each species, we recorded the class, order and family information. The common names of the species were recorded in English and vernacular languages and standardized using the ISO 639-3:2007 code. Besides, information on growth habits, lifecycle (perennial, annual, biennial) and group (monocot/dicot) was also collected for each species.

#### 2.2.2 Pathway and time of introduction

Information regarding the introduction pathway and time was retrieved from the literature reports as well as from global databases. If the introduction pathway of the species could not be identified for India, possible purposes for which the concerned species had spread outside of its native range were identified by searching authoritative literature records. The introduction pathways were categorized following the classification framework proposed by the Convention on Biological Diversity (Harrower et al., 2018). The residence status of a species in India was determined by considering the earliest year recorded in the curated data.

#### 2.2.3 Biogeography

Native and naturalized range information was collated from the global databases at Level 2 (continental and regional scheme) and Level 4 (constituent political units of a country), respectively, as proposed by the International Working Group on Taxonomic Databases for Plant Sciences (TDWG) (Brummitt et al., 2001).

#### 2.2.4 Uses and market dynamics

Usage information of each species was extracted from ethnobotanical literature and databases available in India. The data were supplemented with the usage information reported across the distribution range of a species and standardized according to the thirteen level 1 states of plant uses (Cook, 1995). Data on species trading, including type, quantity and price, for ornamental purposes were collected from e-commerce platforms.

#### 2.2.5 Indian distribution

Occurrence records of the species, downloaded from GBIF (https://doi.org/10.15468/dl.7bkqza), were used to determine their distribution across different states and union territories, land-use and land-cover (LULC) classes, ecoregions, anthropogenic biomes and climate classes in India. LULC information for each of the occurrence records was extracted from the classified image of the country (Reddy et al., 2015). Species’ occurrences were further determined across 51 ecoregions (Olson et al., 2001) and 18 anthropogenic biomes (Ellis and Ramankutty, 2008) present in India. The habitat information at coarser and finer sub-system levels was collected from literature and global databases. The realized climatic niche of each species was characterized for annual mean temperature and precipitation, and their occupancy of the Köppen-Geiger climate classes was ascertained using the *kgc* package (Bryant et al., 2017) in R.

### 2.3 Assessment of data quality

The data quality was assessed based on reliability, usability, and accessibility. The individual data records of the dataset were extracted from peer-reviewed articles and databases that maintain detailed profiles of a large number of species, are frequently updated, and provide traceable data sources. Therefore, the curated data should be considered reliable and consistent. We further adopted a multistep pipeline for technical validation of the curated data. This included random and iterative data checking by individual team members to minimize curator’s bias, avoid omission error and increase the data accuracy before inclusion in the database. To ensure smooth usability of the database, we arranged individual variables in separate .csv files with consistent species names across the data files. The data files were made accessible without any restriction of use through an online data repository and a dedicated website (see ‘Data availability’). A RShiny application (Chang et al., 2020) was created (and embedded on the website) for easy retrieval of species- or variable-specific data through query-based search of the database (see details in Supporting information; Fig.2 shows an example search result).

**Fig. 2:**
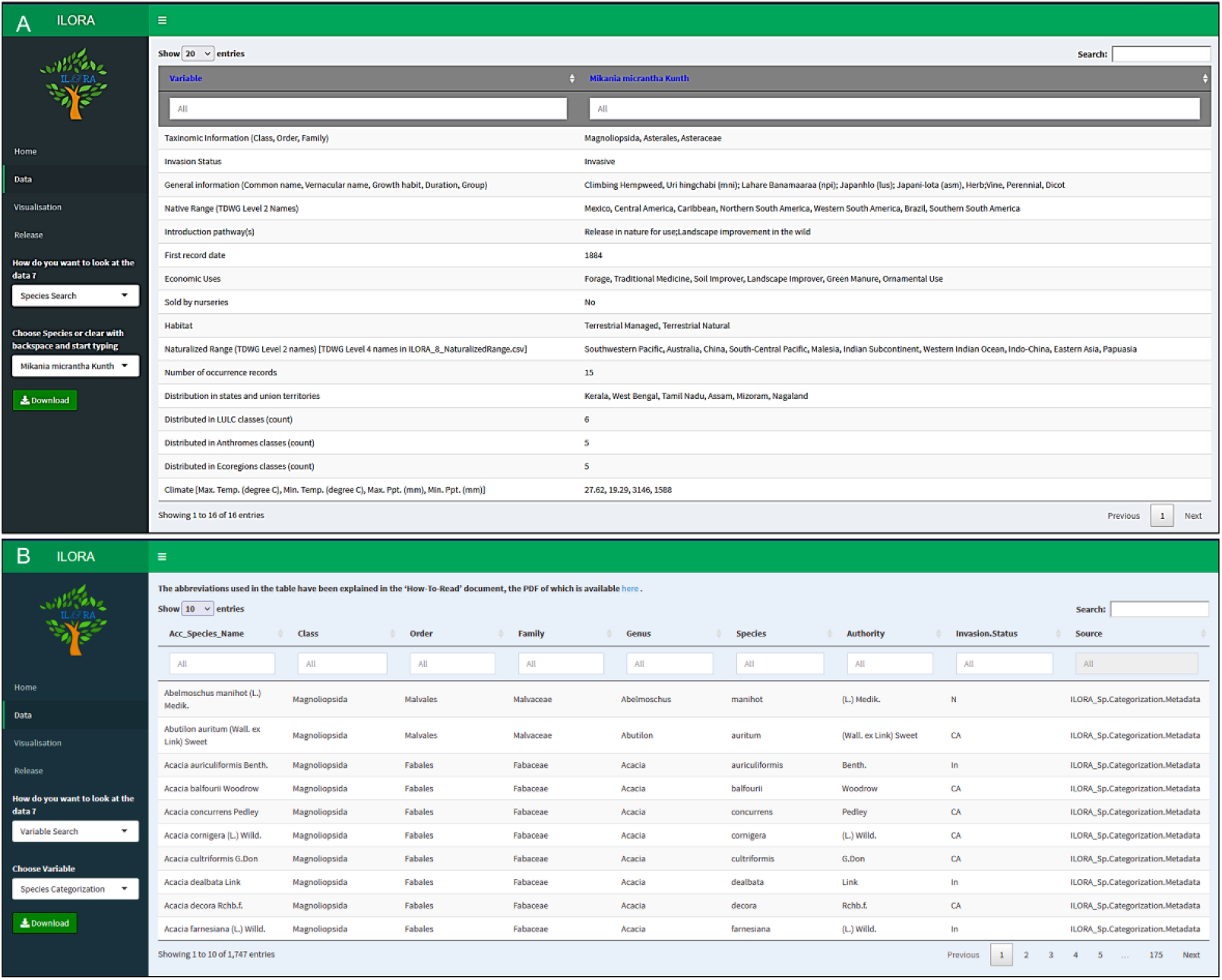
A snapshot of the database search result for – (A) an individual species (*Mikania micrantha* Kunth, as an example) and (B) a variable (Species categorization). The variables for species-specific search (A) follow the arrangement of the data records in the database; note the taxonomic information are displayed separately from the invasion status, as the introduction pathway(s) and the first record date, thus making the number of variables equal to 16

## 3. Usage notes

The primary objective behind curating ILORA is to provide a comprehensive, easy-to-access, policy, and research-relevant database for alien vascular plant species of India. It contains occurrence data and the alien status of a species, both of which were considered as essential variables for invasion monitoring (Latombe et al., 2017). These records can be incredibly useful for stakeholders working on alien plant species in India. It also allows users to comprehend the socio-economic drivers of alien plant spread in the country and formulate appropriate trading policies and quarantine measures for the prevention of future expansion.

Alien species databases are not complete and stable units; dynamism lies at the core of the data records and thus, they are open to suggestions and future updates. ILORA also has provisions for the incorporation of data submitted by the users like researchers, stakeholders and the general public. The submitted data will undergo thorough quality checks by the curators (and domain experts as and when required) before integration into the database. All user-submitted data will accompany source information to ensure full credit to the content providers. This bidirectional information exchange is envisaged to increase the data resolution and would broaden the scope of the database.

## 4. General patterns

Curating data from 22 sources, ILORA version1.0 incorporates attributes of 1747 alien vascular plant species of India in thirteen data variables. Of these, two variables, namely the species categorization and general information, contain data for all the species (Fig.3A). The database contains the recorded common names of 1575 species and vernacular names of 583 species in 23 languages. For the remaining 11 data variables, 1671 species have information for one or more variables whereas data are missing for 76 species (Fig.3B). Among the three categories of alien flora (invasive, naturalized, and casual, Fig.3C), average data availability is maximum for invasive aliens followed by naturalized and casual aliens (Fig.3D). At the taxonomic level, the alien species (invasive, naturalized, and casual) data are distributed across 764 genera and 154 families with Asteraceae, Fabaceae and Poaceae as the most represented families. Out of the 764 genera, 530 genera have species belonging to either one of the three categories (invasive = 76, naturalized = 72, casual = 481) whereas 21 genera have species of all three categories.

**Fig 3.**
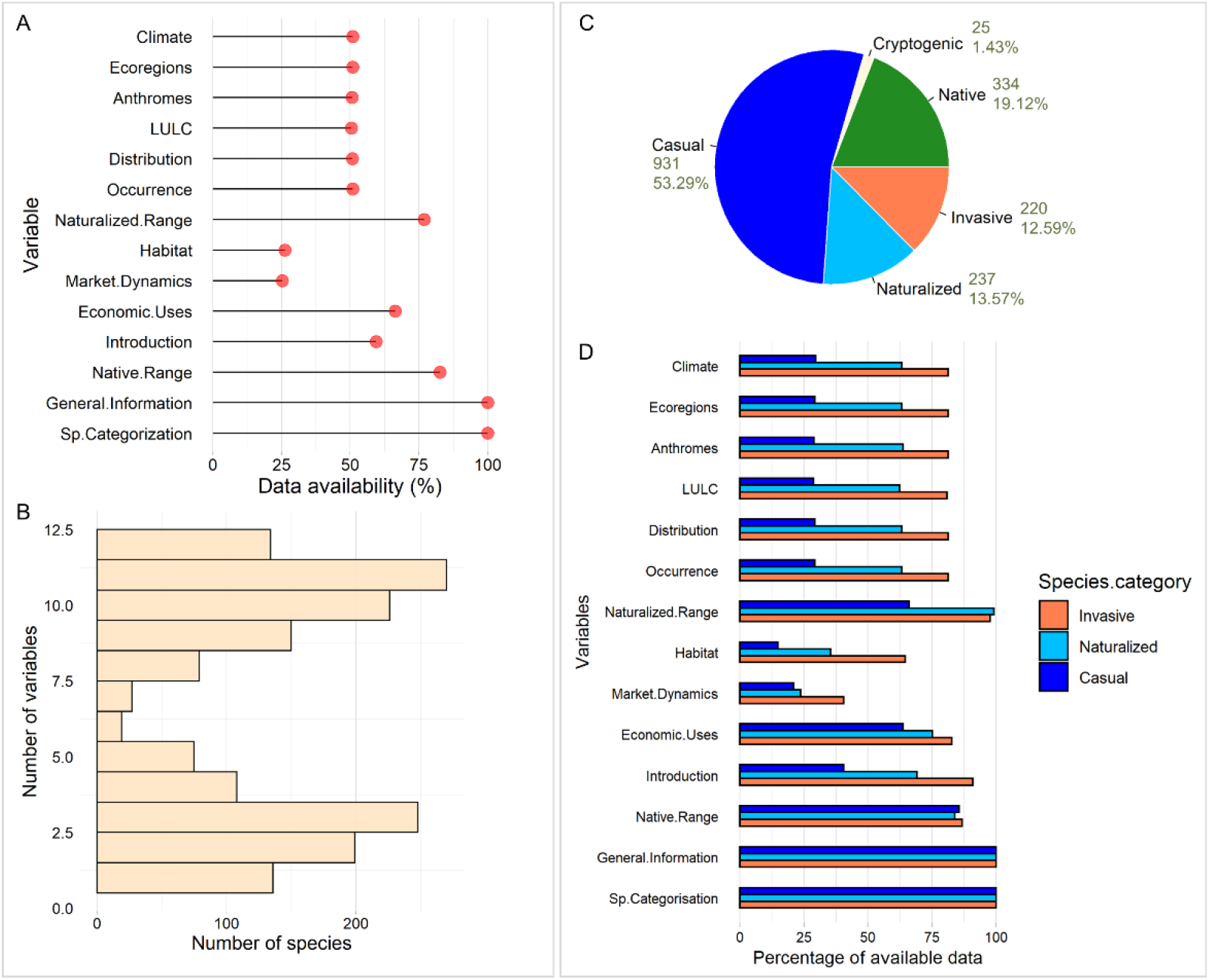
Data availability in the ILORA database – (A) percentage of available data for each of the 14 variables; (B) the frequency histogram showing numbers of species (n=1671 having data for at least one variable) for each of the 12 variables (i.e., excluding two variables, species categorization and general information having no missing data); (C) the pie chart showing the numbers and percentages of the alien flora across five categories; (D) the bar plot showing the percentage of available data for each of the 14 variables for three categories of alien species (invasive, naturalized and casual)

The data records for the species identify 50 native ranges at TDWG Level 2 for 1444 species and 919 naturalized ranges at TDWG Level 4 for 1344 species. Introduction pathway information was available for 1038 species. Available information indicated that the Americas contributed most of the alien species and introduction for ornamental purposes was found to be the dominant introduction pathway (Fig.4).

**Fig.4.**
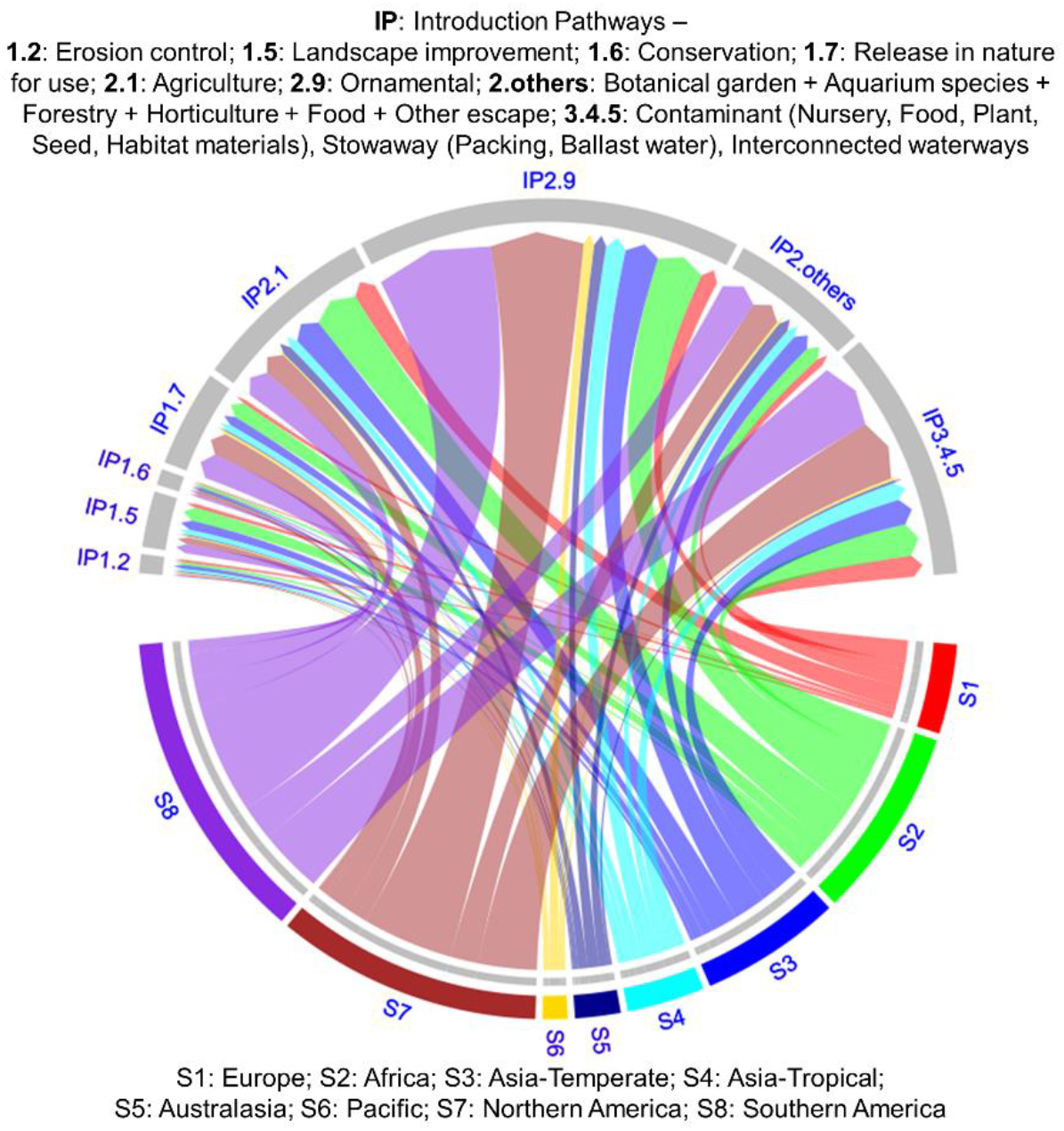
Flows of alien species between native range and introduction pathways. The chord diagram shows the number of alien species (n=317, considering invasive, naturalized and casual alien species for which both native range and introduction pathway information are available in the database) introduced from each continent by different introduction pathways. The width of a chord represents the number of alien species

Herbaceous growth form was predominant across all five categories of species (Fig.5A). The data on potential habitats of 459 species identifies 38 subcategories that are combined into five coarser levels. The database is also populated with 40,724 occurrence records for 889 species, recorded across 34 Land-Use and Land-Cover (LULC) classes, 18 anthromes, 42 ecoregions and 15 climate classes in India. The invasive aliens were found to occupy the maximum number of LULC classes, anthromes and ecoregions (Fig.5B). Among the 36 states and union territories, Tamil Nadu has the highest number of alien flora. Further, 50 subcategories of uses have also been recorded for 1159 species, which were combined into thirteen Level 1 uses (Fig.5C). The majority of species have environmental uses (Fig.5D). The online nursery data for 442 species revealed a higher average market price of native and cryptogenic aliens compared to that of the invasive, naturalized, and casual aliens (Fig.5E).

**Fig.5.**
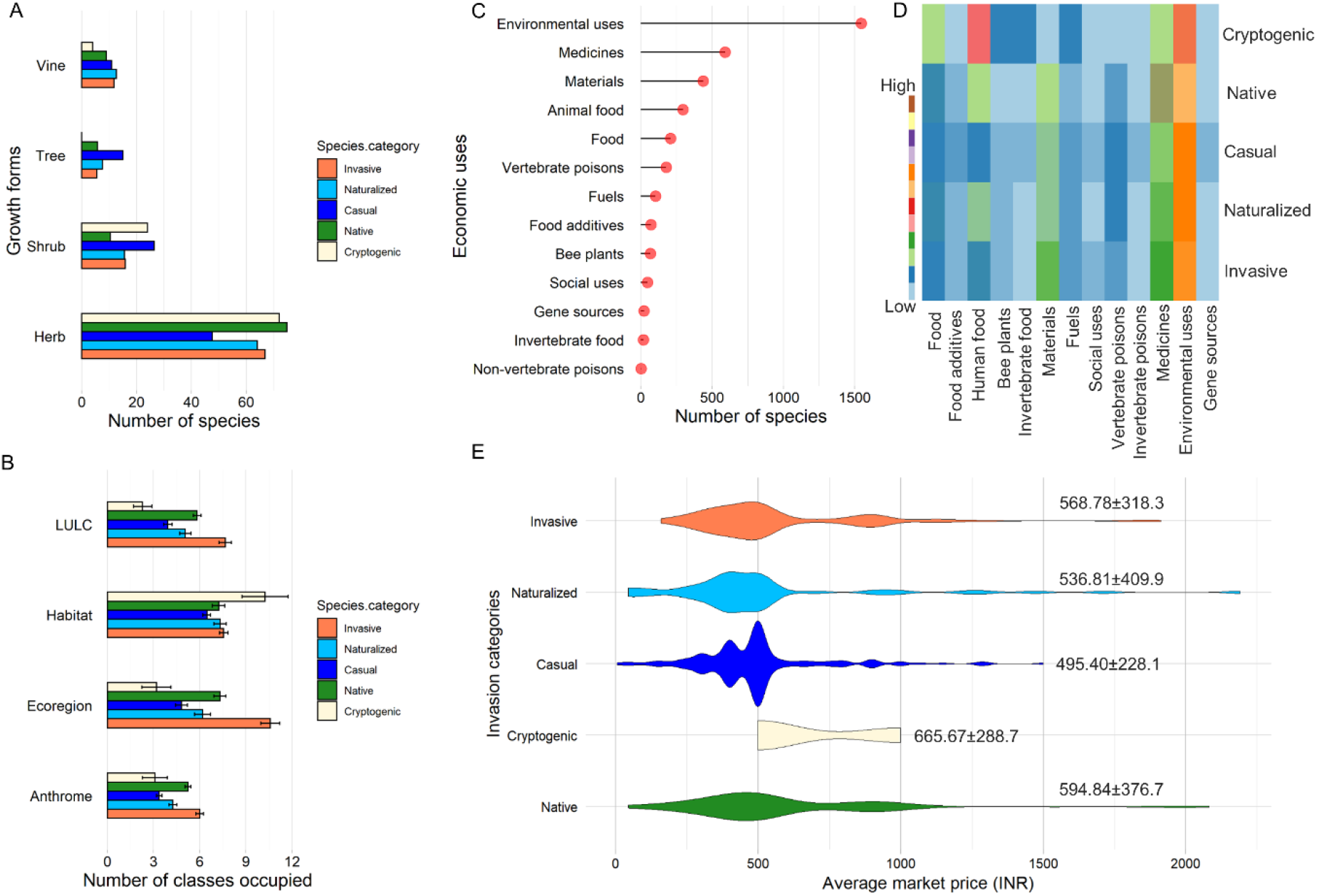
Visualization of variables across five categories of species included in the database – (A) growth form, (B) average (±SE) number of occupied anthromes, ecoregions, habitats and LULC classes, (C) number of economic uses for five categories together, (D) heatmap showing number of economic uses for each of the five categories of species, (E) violin plot showing the distribution of market price values (with mean ± SE values showing above each plot)

## 5. Related works

The introduction pathway and usage data have previously been utilised to propose a set of policy interventions to prevent and control plant invasions in India (Banerjee et al., 2021a). Considering a subset of species, another study identified the variables’ influence on naturalization and invasion success of alien plants in India (Banerjee et al., 2021b).

## Author contributions

AKB and AM conceptualized the idea; VP and AKB developed the methodology; VP, CP and AKB curated the data; all authors validated the data; CP, KP and ARB analysed the data; AKB supervised the project; VP wrote the original draft, which was reviewed by all authors.

## Data availability

The data records, along with the detailed information files and R scripts, are archived in the online repository Figshare (DOI: 10.6084/m9.figshare.13677460). ILORA website (https://ilora2020.wixsite.com/ilora2020) can also be used to retrieve data records through query-based search.

## Acknowledgements

We express our sincere gratitude to our respective institutions and universities for giving us the logistic support, which was an essential requirement in the absence of any specific grant for conducting this research. We are thankful to the associate editor and two reviewers (Prof. Marcel Rejmánek and one anonymous reviewer) for their valuable comments, which have greatly improved the manuscript and database quality. We would also like to thank Prof. Mark van Kleunen, Prof. John Ross Wilson and Dr. Bharat Babu Shrestha, for their thoughtful comments on an earlier version of the manuscript and database.

