## Supporting information for "ILORA: A database of alien vascular flora of India"

**Supporting information**: Methodological details of data curation, assessment of data quality, and data usage and availability

ILORA version 1.0 incorporates attributes of 1747 alien vascular plant species of India for 14 data variables. This document contains details of -

1. Species categorization
2. Rationale for variable selection and data curation
3. Assessment of data quality
4. Data usage and availability
5. Qualitative and quantitative enhancements of the database

### Categorization of the alien flora of India

We developed a multistep pipeline for categorization of alien plant species of India. In the first step, three nationwide checklists, viz. Alien flora of India (1599 species; (Khuroo et al., 2012)), Naturalized alien flora of the Indian states (499 species; (Inderjit et al., 2018)), and the GRIIS- India Version 1.3 (2082 species; (Sankaran et al., 2020)), were consulted to create a comprehensive list of alien plant species of India (n=4180). To remove ambiguity and orthographical errors in the plant nomenclature, automated standardization of taxonomic names was conducted using the *WorldFlora* package (Kindt, 2020) in R version 4.0.2 (R Core Team, 2020) (hereafter, R). This package validates the species names against a static version of the World Flora Online (WFO) taxonomic backbone data, which is actively curated by global experts based on The PlantList backbone (http://www.theplantlist.org/) (static since 2013), and therefore provides the most updated and comprehensive taxonomic reference of vascular plants (Borsch et al., 2020). The list was further cleaned by removing duplicates and synonyms, and the infraspecific taxa and artificially hybridized species were not considered further (n=1677).

The selected plants (n=2503) were subjected to a two-step verification process involving national-level checklists and global databases to determine their – i) origin status (i.e., is the species native or alien to India) and ii) invasion status (i.e., what is the degree of naturalization of the species) (Pyšek et al., 2004) in India. To designate the species categories, we followed the Darwin Core (dwc) standard and adopted the proposed controlled vocabulary for dwc:establishmentMeans (Groom et al., 2019) to maintain stronger links to existing terms and ensure future interoperability across multiple databases. First, the cultivated aliens were identified from the Alien Flora of India (n=777). Among these, the species which were identified as naturalized and/or invasive in either of the four databases, namely the Naturalized alien flora of the Indian states databases, GloNAF, GRIIS and the Germplasm Resource Information Network (hereafter, GRIN: https://www.ars-grin.gov; accessed on 4 April 2020), were removed from the list of the cultivated species (n=21). The cultivated aliens thus identified (n=756) were retained in this version; however, no variable information was extracted for these species.

The origin status of the remaining species (n=1747) was ascertained using three global databases, namely, POWO, CABI-ISC and GRIN (accessed on 20 April 2020). These databases were chosen because they maintain detailed profiles of a large number of plant species which are frequently updated, provide traceable data sources, and are widely referred by plant biologists to trace the origin of a species (Martin et al., 2017; Turbelin et al., 2017). If the origin status of a species was identified as 'native' or 'alien' to India in at least one of these databases and was not contradicted by the remaining (n=1576), it was accepted. However, in case of any contradiction between databases (e.g., two databases identified a species as alien whereas one identified it as native) (n=162), peer-reviewed journal articles, published books, and online databases were consulted to establish the origin status of the species. For literature and book search, the Google Scholar database (https://scholar.google.com/; accessed on 29 April 2020) was used. The search terms were optimized following a preliminary search based on a handful of relevant articles retrieved by using the names of the species. The optimized search string consists of a combination of species name and nativity (and its synonyms and related words). The advanced search function of the Google Scholar database was used to search for "species name" with at least one of the words "native" OR "endemic" OR "native region" OR "nativity" OR "origin" present anywhere in the article. The search results were screened until the origin status of the species was discerned confidently. The online databases and repositories were retrieved through a Google search (http://www.google.com/; accessed on 29 April 2020) and the databases were considered based on the same criteria (i.e., frequently updated and with traceable data sources). The species with ambiguous origin status (i.e., which cannot be resolved by the global databases and literature search) were categorized as 'cryptogenic' (i.e., uncertain biogeographic status) following (Essl et al., 2018). At the end of this step, the origin status of the alien species was categorized as – alien (n=1388), native (n=334) and cryptogenic (n=25).

In the second step, the status of the alien species in the introduction-naturalization-invasion continuum was determined. From the list of alien species (n=1388), the 'invasive' species were categorized by consulting the GRIIS database (i.e., the species identified as 'Alien Invasive' in GRIIS) (n=220) as it provides the latest list of invasive alien species present in India. The species which have been identified as naturalized aliens by the Naturalized Alien Flora checklist and the GloNAF database, were categorized as 'naturalized' (involving both invasive and non-invasive naturalized ones) (n=237). The remaining alien species were identified as 'casual alien' following (Blackburn et al., 2011) (n=931). Finally, at the end of this step, the assembled list of alien flora of India was classified into six categories, namely invasive, naturalized, casual alien, cryptogenic, native, and cultivated species. We included the native species in this database version because these species were identified as aliens in one of the three national checklists and expert in-depth knowledge in future may provide the conclusive status of these species (see 'Scopes of enhancements' section).

It is important to note here that assigning species categories to alien/native (origin status) and invasive/naturalized/casual (invasion status) based on detail biogeographic information is out of scope of ILORA version 1.0. Rather, we intend to present the available information for these species across national and global databases and identify their status in India at a certain confidence level (Table 1 and Table 2). In absence of adequate biogeographic knowledge for a large number of species in India, this approach provides a broad idea of species status in India. The framework used here is open for future development so that new information about species status can be accommodated when available. Nevertheless, the species status reported here should bridge the knowledge gaps, assist further research and most importantly, ignite scientific discussion on ambiguous species status.

Table 1: Logic for assigning confidence score (0-1) to origin status of alien species based on their categorization by the national and global databases

| Database | Possible combination | | Confidence score | No. of species |
|---|---|---|---|---|
| Agreement across databases | | | | |
| Any 1 | Two | 1 India-specific + 0 general | 0.65 | 126 |
| | | 0 India-specific + 1 general | 0.6 | 106 |
| Any 2 | Three | 2 India-specific + 0 general | 0.8 | 237 |
| | | 1 India-specific + 1 general | 0.75 | 269 |
| | | 1 India-specific + 0 general | 0.7 | 473 |
| Any 3 | Four | 3 India-specific + 0 general | 1.0 | 131 |
| | | 2 India-specific + 1 general | 0.95 | 118 |
| | | 1 India-specific + 2 general | 0.9 | 69 |
| | | 0 India-specific + 3 general | 0.85 | 47 |
| Disagreement between databases – External search supports | | | | |
| Any 1 | Two | 1 India-specific + 0 general | 0.4 | 60 |
| | | 0 India-specific + 1 general | 0.35 | 51 |
| Any 2 | Three | 2 India-specific + 0 general | 0.55 | 18 |
| | | 1 India-specific + 1 general | 0.5 | 11 |
| | | 1 India-specific + 0 general | 0.45 | 6 |

Note: 'India-specific' denotes native or alien information available in the consulted databases are specific to India; 'General' means otherwise

Table 2: Logic for assigning confidence score (0-1) to invasion status of alien species based on their categorization by the national and global databases

| | | | | |
|---|---|---|---|---|
| When Category = "Invasive" | | | | |
| AFI | NAFIS | GLONAF | Score | No. of species |
| Invasive, Native/Invasive, Naturalized | Naturalized | Naturalized | 0.95 | 141 |
| Invasive, Native/Invasive, Naturalized | Blank | Naturalized | 0.9 | 17 |
| Invasive, Native/Invasive, Naturalized | Naturalized | Blank | 0.9 | 7 |
| Invasive, Native/Invasive, Naturalized | Blank | Blank | 0.8 | 14 |
| Others | Naturalized | Naturalized | 0.9 | 8 |
| Others | Blank | Blank | 0.7 | 32 |
| Others | Naturalized | Blank | 0.85 | 1 |
| Others | Blank | Naturalized | 0.85 | 0 |
| When Category = "Naturalized" | | | | |
| AFI | GRIIS | | Score | No. of species |
| Invasive, Native/Invasive, Naturalized | Alien | | 0.95 | 129 |
| Invasive, Native/Invasive, Naturalized | Blank | | 0.9 | 22 |
| Others | Alien | | 0.9 | 66 |
| Others | Blank | | 0.8 | 20 |
| When Category = "Casual Alien" | | | | |

| AFI | GRIIS | Score | No. of species |
|---|---|---|---|
| Not blank | Alien | 0.95 | 142 |
| Not blank | Blank | 0.8 | 18 |
| Blank | Alien | 0.9 | 771 |
| Blank | Blank | 0.7 | 0 |
| When Category = “Cryptogenic” | | | |
| AFI | GRIIS | Score | No. of species |
| Blank | Blank | 0.95 | 3 |
| Blank | Alien | 0.9 | 14 |
| Not blank | Alien | 0.8 | 6 |
| Not blank | Blank | 0.9 | 2 |
| When Category = “Native” | | | |
| AFI NAFIS GLONAF GRIIS | | Score | No. of species |
| All 4 = Not blank | | 0.7 | 18 |
| Any Three = Blank | | 0.9 | 86 |
| Any Two = Blank | | 0.8 | 94 |
| Any One = Blank | | 0.75 | 136 |
| All four = Blank | | 0.95 | 0 |

Note: AFI – Alien Flora of India (Khuroo et al., 2012); NAFIS: Naturalized Alien Flora of the Indian States (Inderjit et al., 2018); GLONAF: The Global Naturalized Alien Flora database (van Kleunen et al., 2019); GRIIS: Global Register of Invasive and Introduced Species (Sankaran et al., 2020)

### Data curation

#### Taxonomy and general information

Taxonomic classification is essential for identification and analysis of evolutionary relationships between the species and hence, is an indispensable part of any plant-based database. The taxonomic information (class and order) was recorded from POWO (and from CABI-ISC, if not available in POWO) and the genus, species and family names were retained as obtained from the standardization of species names using the WFO taxonomic backbone. The phylogenetic relationship between the three categories of alien species (invasive, naturalized and casual) was resolved through 'mega-tree' approach by using the *V.PhyloMaker* package (Jin and Qian, 2019) in R (specifically, using the options of build.nodes.1 and scenario 3). The common names of the species (in English) were documented from GRIN, POWO and CABI-ISC databases. Since, the vernacular names are easier to relate to and may increase the accessibility and usability of the database, we curated the vernacular names of the species from two online repositories, namely the eFlora of India (https://efloraofindia.com/; accessed on 31 May 2020) and Flowers of India (http://flowersofindia.net/; accessed on 31 May 2020), and the digital database of Flora of Peninsular India (http://flora-peninsula-indica.ces.iisc.ac.in/welcome.php; accessed on 31 May 2020). The vernacular names were standardized using the ISO 639-3:2007 code consisting of three-letter language identifiers for the representation of languages.

Along with taxonomic identity, the general information of each species, namely its growth habit, duration (perennial, annual, biennial) and group (monocot/dicot), was collated from the TRY Database (Kattge et al., 2020) (accessed on 23 May 2020), a comprehensive web-archive that integrates nearly 400 different plant traits. If there was missing data in TRY, databases focused on Indian flora, namely the digital databases of Flora of Peninsular India (http://flora-peninsula-indica.ces.iisc.ac.in/welcome.php; accessed on 31 May 2020), India Biodiversity Portal and Flowers of India (http://flowersofindia.net/; accessed on 31 May 2020), were checked to extract general information of the species. Further, global databases like the database of the Natural Resources Conservation Service (NRCS) (https://www.nrcs.usda.gov/wps/portal/nrcs/site/national/home/; accessed on 28 May 2020) was used to obtain information. For the small number of species, for which data was absent in all of the above databases, Google search engine (https://www.google.com/; accessed on 31 May 2020) was used, and relevant information was retrieved using the following keyword combinations: species name, growth habit and duration (including all categories of growth habit and duration).

#### Pathway and time of introduction

Residence time is an important predictor of naturalization; the longer a species is present in a region, the more propagules it can produce, resulting in a higher probability of founding new populations (Křivánek et al., 2006). A positive relationship is, therefore, frequently found between the residence time in an introduced range and the naturalization success of an alien species (Feng et al., 2016). It is therefore not surprising that identification and prioritization of the introduction pathway of alien invasive species have been recognized as one of the Aichi Biodiversity Targets (Target 9) (www.cbd.int/decisions/cop/?m=cop-10; accessed on 1 June 2020) and retained as one of the targets (Target 5) to be achieved by 2030 in the zero draft of the post-2020 global biodiversity framework (https://www.cbd.int/doc/c/3064/749a/0f65ac7f9def86707f4eaefa/post2020-prep-02-01-en.pdf; accessed on 13 September 2020).

Information regarding the introduction pathway was retrieved from Google Scholar. We used the advanced search function of the Google Scholar database (https://scholar.google.com/#d=gs_asd;

accessed on 20 May 2020) to search for a species name with at least one of the search queries related to introduction purpose ["introduction history" OR "introduced" OR "introduction purpose" OR "introduction pathway"] present anywhere in the article with the exact phrase “India”. The screening of articles was continued until the relevant information for a specific introduction purpose was obtained for a species. In addition to these sources, we relied on our knowledge of the Indian literature and searched two other journals, namely Tropical Ecology archives and Indian Forester, which are not indexed in the Google Scholar database. Information for the remaining species (for which there was no data available from the literature) was retrieved from CABI-ISC database (accessed on 12 June 2020). If the introduction purpose of the species could not be established for India, then possible purposes for which the concerned species had spread outside of its native range were identified by searching authoritative literature records. A structured query search string ["species name" AND ("introduction history" OR "introduction pathway" OR "introduction purpose" OR "introduced")] was used to retrieve this information using ISI Web of Science platform (https://webofknowledge.com/; accessed on 13 June 2020). The reasons for introductions were categorized further following the introduction pathway classification framework proposed by the Convention on Biological Diversity (Harrower et al., 2018).

We recorded the introduction time of the species mentioned in the literature. In addition, the oldest occurrence record of the species in the country was also retrieved from – 1) the Global Biodiversity Information Facility (GBIF) (http://www.gbif.org; accessed on 16 June 2020) and 2) the Global Alien Species First Record Database version 1.2 (Seebens et al., 2017). The GBIF database was suitable for this exercise due to its comprehensive wealth of information on museum specimens collected in the 18$^{th}$ and 19$^{th}$ century whereas the Global Alien Species First Record Database holds region specific first records of 47542 alien established taxa. To assess the residence status (sensu (Pyšek et al., 2004)) of the species in India, we considered the earliest year recorded among literature and these two databases. Finally, the information on the native range for each of these species was retrieved from GRIN database (accessed on 4 April 2020) following the continental and regional scheme (Level 2) proposed by the International Working Group on Taxonomic Databases for Plant Sciences (TDWG) (Brummitt et al., 2001).

### Biogeography

More than 13,000 vascular plant species have already become naturalized outside of their native ranges (van Kleunen et al., 2015), and the naturalization risk of alien species is predicted to increase due to climate change (Dullinger et al., 2017). Further, a recent study has reported that warm temperate and sub-tropical regions have a higher relative richness of naturalized alien plant species (Essl et al., 2019). In this context, information on the geographical range, where each of the species has become naturalized across 1029 global regions, was recorded from the GloNAF database. The naturalized ranges of the species were recorded by identifying the subset of geographical areas that fall within the political boundary of India (region_id: 380-402 in the GloNAF database). This provided us with the information on global as well local (nationwide) naturalized ranges for each of the species. The finest resolution of naturalized range information provided by GloNAF is TDWG-Level 4, corresponding to the constituent political units of a country, i.e., states and union territories of India. The TDWG-Level 4 range information for the other species (i.e., except the naturalized ones) was retrieved from the CABI-ISC database (accessed on 12 June 2020).

### Uses and market dynamics

Economic activities have the potential of acting as direct (Dehnen-Schmutz et al., 2007; Faulkner et al., 2016) as well as indirect drivers (Liu et al., 2019; Ward et al., 2020) of plant invasion. The synchronous increase in trade, transport, and tourism act as a major driving force behind the increasing alien species introductions across the globe (Anderson et al., 2015; Hulme, 2009). Additionally, most of the intentionally introduced plant species tend to have multiple economic uses, e.g., cultivation, ornamental, medicinal uses, which can increase the propagule pressure and promote naturalization success (van Kleunen et al., 2020). To capture such economic usage information for the alien flora of India, we first extracted information from the book Ethnobotany of India (Pullaiah et al., 2016). The five volumes of this book provided detailed and updated information of plant uses focusing on major biogeographical zones of the country. The medicinal use in India was recorded in binary format (Present/Absent) by surveying the online database of the Environmental Information System's Centre on Medicinal Plants (http://envis.frlht.org/; accessed on 14 June 2020), operational under the aegis of the Ministry of Environment, Forests and Climate Change (MoEFCC), Government of India. Two additional databases, namely GRIN and CABI-ISC (accessed on 12 June 2020), were also explored. These databases provided information on economic uses reported across the distribution range of each species. Also, Useful Tropical Plants (http://tropical.theferns.info/; accessed on 15 June 2020) and Useful Temperate Plants (http://temperate.theferns.info/; accessed on 15 June 2020) databases were probed to document the ornamental uses. The thirteen level 1 states of plant uses as prescribed by TDWG in their Economic Botany Data Collection Standard (Cook, 1995) was then used as a template to categorize the uses for each of the species.

Ornamental trading of species is known as one of the major drivers for the introduction and spread of alien plant species worldwide (Dehnen-Schmutz et al., 2007; Niemiera and Von Holle, 2009; van Kleunen et al., 2018). To comprehend the trading scenarios of these species in India for ornamental purposes and their corresponding translation to revenue streams, two e-commerce platforms were scouted. In India, the e-commerce industry has been on an upward growth trajectory and is recognized as the fastest growing industry in India (https://www.ibef.org/industry/ecommerce.aspx, accessed on 10 July 2020). Further, e-commerce trade, being devoid of strict bio-security regulations, is providing a new distribution channel for alien and invasive species worldwide and India is no exception (Humair et al., 2015). For this exercise, we considered two online nurseries, namely Nurserylive (https://nurserylive.com) and Plantslive (https://www.plantslive.in), as the representatives of online ornamental plant trading scenario in India because they – i) mention sales history of each plant species, ii) provide national-level shipping facilities, and iii) include the scientific names of each species in their record. Data collected from these portals included types of plant material being sold (seed/plant), quantities of each species sold during the previous month (i.e., June 2020; retrieved on 15 July 2020), and their unit price (in INR).

### Indian distribution

Information on the current distribution of alien species in a geographical landscape is important to understand the ecological and environmental interactions shaping the introduction to naturalization continuum and has a broad range of applications regarding the management of naturalized and invasive species (Zhou et al., 2020). To estimate the geographical distribution of the species across the country, their occurrence records (latitude and longitude) species were extracted from GBIF (accessed on 5 July 2020; https://doi.org/10.15468/dl.7bkqza). Since the web interface allows downloading of occurrence data for a maximum of 200 species, we used the *rgbif* (Chamberlain et al., 2016) and *taxize* (Chamberlain and Szöcs, 2013) packages in R to search and download the occurrence records by matching the taxon keys with GBIF and by using the following predictors: country (India), coordinate (yes), geospatial issue (false). The GBIF database was searched with the standardized taxon names along with their corresponding author names (see 'Categorization of the alien flora of India' section). The occurrence records were cleaned thereafter to align the data with the mother database. First, the points falling in the ocean were removed from the dataset. Second, as GBIF retrieves occurrences using synonymous species names, the occurrences of the synonymous species were merged. Similarly, occurrence records were merged for the species that had more than one author names, and the author's name was retained according to the mother database. Finally, the occurrence data of the infraspecific taxa were removed to align the occurrence dataset with the species-level mother database. The occurrence records were then used to determine the distribution of each of the species in the states and Union Territories (UTs) of the country. The occurrence records were intersected with the state and UTs' boundaries by using the intersection tool in ArcMap 10.2.1. The resulting information was further processed using a custom script in R to identify the presence or absence of the species in the states and UTs of India. The occurrence records were further used to determine the species' presence across different land-use and land-cover (LULC) classes, ecoregions, and anthropogenic biomes present within the geographical boundary of the country.

To identify the species' distribution across the different LULC classes, we used a recently classified image of the country having a spatial resolution of 56 m acquired by the multi-season IRS Resourcesat-2 Advanced Wide Field Sensor (AWiFS) (Reddy et al., 2015). The classified image identified land-use and land-cover classes based on natural vegetation types (excluding cultivated systems, croplands, plantations, barren land, water bodies, and snow). By using the spatial extraction tools in ArcMap 10.2.1, LULC information for each of the occurrence records was extracted. This information was then processed in R to identify the presence or absence of the species in 34 LULC classes. Notably, LULC is a dynamic variable and information of LULC classes occupied by a species should be treated as the potential LULC classes which the species can occupy and in no way reflects its current distribution.

The species' occurrences were further determined across different ecoregions and anthropogenic biomes. An ecoregion is characterized as land having a distinct assemblage of species and natural communities with boundaries *a priori* of major land-use change (Olson et al., 2001). The anthropogenic biomes, also known as anthromes, characterize anthropogenically-altered terrestrial biosphere by using patterns generated from human-ecosystem interactions (Ellis and Ramankutty, 2008). Out of the 814 global ecoregions, 51 were found in India distributed within 12 biomes and the Indo-Malay biogeographic realm. Out of the 21 global anthromes, 18 anthromes were found in India. By using the intersection and spatial extraction tools in ArcMap 10.2.1, we identified the species' distribution across the different ecoregions and anthromes present in the country using a custom script in R.

In addition, the information regarding the habitats of the species was also collected from CABI-ISC (accessed on 12 June 2020). The habitat information was extracted at two levels: the coarser system level (terrestrial, freshwater, littoral and brackish) and the finer sub-system level (e.g., forests, riverbanks, wetlands, deserts, urban areas).

The bioclimatic variables, derived from the monthly temperature and rainfall values, are considered as biologically meaningful variables and are used in various ecological modelling studies (Jeschke and Strayer, 2008). Occurrence records were used to characterize the realized climatic niche of the species for two bioclimatic variables - annual mean temperature (BIO1) and annual precipitation (BIO12). The bioclimatic variables (averaged over 1970-2000) were downloaded from WorldClim version 2 at a spatial resolution of 2.5 arc minutes (approximately 5 km resolution at the equator) (Fick and Hijmans, 2017). The BIO1 and BIO12 values were extracted for each of the occurrence records using the Spatial Analyst Toolbox of ArcMap 10.2.1. The extracted information was further processed in R to identify the range (minimum and maximum) and average (±SD) annual temperature and precipitation values for each species. Notably, temperature and precipitation values should be interpreted as realized niche of a species (i.e., from where the species has been recorded yet). The fundamental niche of the species is ‘larger’ than the realized niche (Hutchinson, 1957) and can only be detected when the species colonizes new environmental conditions by overcoming competitive interactions and other limiting factors which are responsible for restricting the realized niche of the species (Soberón and Arroyo-Peña, 2017).

The distribution of the species across different climatic zones in India was further identified. For this purpose, we used the Köppen-Geiger empirical climate classification system, which classifies the global climate into five major classes and 30 sub-types (Köppen, 1936). The species' occupancy of the climate classes present in India was ascertained using the *kgc* package (Bryant et al., 2017) in R, in which occurrence records were used as input data.

### Assessment of data quality

The data quality was assessed by using a checklist having three criteria – reliability, usability, and accessibility. The individual data records of the dataset were extracted from highly-cited and well-regarded databases, peer-reviewed authoritative scientific journals, and published books. Therefore, the curated data should be considered as reliable and consistent. However, there are many sources of uncertainty in such data (McGeoch et al., 2012). We further adopted a two-step process for technical validation of the curated data. Intense discussions among the developer team were undertaken to maintain the quality of the adopted strategies and techniques of data collection. First, the citations and sources provided by the larger databases were randomly checked to acknowledge the quality of the data provided. Hence, the accuracy of the data sources was successfully established. Second, after completion of data curation for each variable, data for 10% species were rechecked by another curator of the team. The mismatches between data collected by two curators, if any, were resolved and the rechecking process was repeated with 15% species. The process was repeated with an increment of 5% species at every turn until there was no variation between curators' observations. Third, duplicate data extraction was performed for 10% species (including species with missing values) to remove exclusion bias (i.e., to avoid omission errors). Finally, before including the data in the mother database, profiles of 10% species (n = 175) were rechecked by an individual team member. At this stage, we noted no mismatch between individual team member's observations, thereby reliability of the curated data was assured.

To ensure smooth usability of the database by the end users, we arranged individual variables in separate .csv files with keeping species names consistent across the data files. We further included a document in the database as a standalone file explaining the column heads for each data file separately. Finally, the data files were made accessible without any restriction of use through an online data repository and a dedicated website, the details of which have been provided in the 'Data accessibility' section.

### Data usage

The primary objective behind curating ILORA is to provide a comprehensive, easy-to-access, policy, and research-relevant database for alien vascular plant species of India. It contains data of two (alien species occurrence and species alien status) of the three Essential Variables for Invasion Monitoring (except the alien species impact) (Latombe et al., 2017) of alien plant species present in India. ILORA should not be considered as a checklist of alien flora present in India. Rather, the essential and the supplementary variables (i.e., the derived variables like the pathways of introduction and characteristics of the receiving environment), which have been collated both *in-situ* and from *ex-situ* sources (Latombe et al., 2017), provide a baseline data that can be utilized for identifying, reporting, monitoring, and regulating the status of alien plant species in India. With data records ranging from taxonomy, perceived alien status, introduction history to socio-economic uses, and geographical distribution, ILORA envisages assisting the various stakeholders who are associated with alien plant invasion research and policymaking in India.

The data records of ILORA could be incredibly useful to cater to the need of data availability for research on alien plant species in India. For example, the database can be used to identify the relative importance of variables along with the progression of alien species through the introduction-naturalization-invasiveness continuum. The database can be of immense value to identify potential invasive species. A recent study has modelled the impacts of the environment, species characteristics and nursery availability to identify the predictors for naturalization and to forecast naturalization risks of ornamental plant species in Europe under future climate conditions (Haeuser et al., 2018). Similar modelling exercises could be crucial in understanding the dynamics of the alien species and in streamlining the management efforts towards timely containment or eradication, especially for a developing country like India. Further, the broad range of distribution information can be used for habitat suitability, correlative and mechanistic distribution modelling approaches for predicting the potential distribution and range expansion of naturalized and invasive species as well as of the emerging invaders in the country. These findings may inform policy directives for timely management of the alien plant species based on their residence status across the invasion growth curve. On the other hand, data on introduction purpose and naturalized range of the species could be used for formulating international trading policies, imposing stricter border control measures to prevent novel introductions, as well as developing domestic trading policy interventions to check the further spread and naturalization of species. ILORA also allows users to comprehend the socio-economic drivers of alien plant spread in the country. Similarly, information on the distribution of species across different climate classes, anthromes, ecoregions, and LULC classes could guide the implementation of site-specific management protocols to prevent naturalization of alien species.

ILORA has also provisions for future users to contribute data and information to ensure bidirectional information exchange. Users can submit data for all 13 variables included in version1 of the database for the species already present in the database and also for new species to be included. The standard for such data submission has been given on the database website. For each variable, users need to provide detail information of the data sources which can be traceable and authenticated. Such sources include, but are not limited to, published journal article (preferably with DOI), book or book sections (preferably with DOI), historical text, thesis report, conference paper, published dataset, online database, and government document. In case the submitted data is not verifiable by an authenticated source (e.g., personal observation), users need to submit detailed descriptions of the data collection (e.g., photographs with date and time stamps with the GPS feature enabled). The submitted data will undergo verification and thorough quality-checks by the curators (and domain experts as and when required) before integration in the database to ensure data quality. All data will be associated with source information and ILORA will encourage users to refer back to the primary sources so that the content providers can get full credit and acknowledgement. This practice will allow expansion of the resolution as well as the scope of the database.

### Data availability

The data records can be accessed through two pathways –

1) Online data repository: The data records are arranged and archived into the 14 files (in .csv format) in the online repository figshare (DOI: 10.6084/m9.figshare.13677460). In each of these files, the records are associated with 'Acc_Species_Name' which allows correspondence across the entire database. The file ILORA_summary.csv contains an overview of the data records available for individual species. Cells with missing or unavailable information are left blank. An overview of the file name, data contents and data type have been provided in Table 1.

2) Dedicated website: ILORA website (https://ilora2020.wixsite.com/ilora2020) is a user-friendly, public platform that can be used to retrieve necessary data records through query-based search of the database. A RShiny (Chang et al., 2020) application has been created and embedded on the website for providing easy and complete access of the database in an interactive manner. Users can search the database by species name or for specific variable. An advanced search function is also present to facilitate data retrieval through multiple combinations of search criteria. The data files or a subset of the data (based on query) can be downloaded in .csv or in plain text format. The scripts used to process the data files are also available in a GitHub repository (https://github.com/Ilora2021/ILORA/). The website is also supplemented with data highlights, visual representation of data, and associated infographics, which enables the information to permeate a broader range of audiences. It also provides a feedback loop for the users to input new data or update the existing data, thereby contributing to the dynamism and long-term sustainability of the database. As is the common practice for any database, updated information will be communicated to the subscribed users on a regular basis.

### Scopes of enhancements

Alien species databases are not complete and stable units; dynamism lies at the core of the data records and thus, they are open to suggestions and future updates. ILORA is also not exhaustive in every aspect as there is a lot of scope for the enrichment of data resolution. The subsequent versions of the database will also increase data resolution by including – i) detail profiles of the cultivated aliens (since many of these species may escape human control and become naturalized; all data files), ii) potential dispersal pathways (valuable for the prevention of further spread and reduce chances of novel invasion; ILORA_Introduction.csv), iii) trading scenarios (both online and offline sales figures of alien plant species in the market which can be used as a proxy for propagule pressure; ILORA_MarketDynamics.csv), and iv) functional traits (following the leaf-height-seed scheme proposed by (Westoby, 1998); ILORA_GeneralInformation.csv).

While we, as the developers, will work towards increasing the quantitative and qualitative enhancements of ILORA, we have also identified the following scopes which grant opportunities for contributions from the scientific community, citizens, and stakeholders for further improvement of the data quality. First, the origin status of some of the alien species was found to be contradictory between the national checklists and global databases. Such discrepancies may arise from knowledge gaps between respective curators leading to lack-of-consensus decisions; nevertheless, it can be resolved by the thorough and scientifically-informed pursuit of the literature and archival records. However, this exercise is often challenging, given that many such records are not systematically digitized or included in searchable databases. One possible way around this is to rely on in-depth biogeographic knowledge accumulated from the working experience of the concerned taxa. Further, given the large geographic extent of the country, it is likely that some species are native to one part of India but naturalized in other parts. For example, *Nasturtium officinale* has been recognized as naturalized in India across the three databases (Inderjit et al., 2018; Khuroo et al., 2012; van Kleunen et al., 2019). According to POWO, the species is native in Pakistan whereas GRIN recognizes the species as native in India. If pre-partitioned British India is considered, the species is likely to be native in northern India (a politically different entity from Pakistan, but biogeographically similar) from where it might have been introduced to peninsular India and became naturalized (personal communication). Therefore, the resolution of origin and invasion status of the alien species can be increased many folds by considering documented introduction history and robust biogeographical evidence, if available, on a case-by-case basis (ILORA_Sp.Categorization.csv). Similarly, the majority of the invasive plant species reported in different countries in South Asia were reported or imported first in India. Initiatives similar to ILORA by other South Asian countries, or preferentially a common database of this kind, will be an important step towards preventing and managing invasive alien species at the regional level. By providing the baseline information available for each species, ILORA opens the door for further discussion among the scientific community and expects knowledge support from the researchers, especially from the taxonomists and phylogeographers, to refine the database.

Second, while the database curated the available information on the introduction pathway at the highest available resolution, we are aware that detailed scientific studies of a particular taxon can generate novel information or modify the existing knowledge. For example, *Mikania micrantha* (Compositae) has been long known to be introduced during the 2$^{nd}$ World War to camouflage the airfields of northeast India (Tripathi et al., 2012). However, reconstruction of the introduction pathways based on herbarium records and archival literature revealed multiple introductions of the species in different regions in India (Banerjee et al., 2019). Similar fine-scale information is only available for a few species, primarily due to focused research interest of Indian academia and unavailability of archival and herbarium records in a digital and easy-to-access format. ILORA, therefore, welcomes the contribution of scientific knowledge for species with unknown introduction pathways and modification of the existing information, if any, by presenting evidence (ILORA_Introduction.csv).

Third, the resolution of the occurrence records as well as of the distribution information can be improved by many folds. Presently, the occurrence records were extracted from GBIF due to its comprehensive coverage of both historical herbarium collections as well as contemporary citizen contributions. However, the occurrence records in GBIF are not exhaustive and often poorly represent the actual distribution of a species. For example, for a widespread species like *M. micrantha*, GBIF has only 15 records from India which can be certainly improved by including personal observations, research data and herbaria information. If collected, this information would help in achieving a higher data resolution, as demonstrated by the FLOTROP database on plant diversity of northern Africa by including locally available information with the existing data in GBIF (Taugourdeau et al., 2019). While we will work towards curating occurrence records from the published literature (ILORA_Occurrence.csv), it may not be possible for us to capture the vast information stored in local herbaria and reports in vernacular languages. The occurrence data stored in our database would surely be valuable for broad-scaled research and applications, and contributions from future users of ILORA of hitherto unavailable information could significantly increase the resolution of the database.

van Kleunen, M., Dawson, W., Essl, F., Pergl, J., Winter, M., Weber, E., Kreft, H., Weigelt, P., Kartesz, J., Nishino, M., 2015. Global exchange and accumulation of non-native plants. Nature.

van Kleunen, M., Essl, F., Pergl, J., Brundu, G., Carboni, M., Dullinger, S., Early, R., González-Moreno, P., Groom, Q.J., Hulme, P.E., Kueffer, C., Kühn, I., Máguas, C., Maurel, N., Novoa, A., Parepa, M., Pyšek, P., Seebens, H., Tanner, R., Touza, J., Verbrugge, L., Weber, E., Dawson, W., Kreft, H., Weigelt, P., Winter, M., Klonner, G., Talluto, M.V., Dehnen-Schmutz, K., 2018. The changing role of ornamental horticulture in alien plant invasions. Biol Rev 93, 1421-1437.

van Kleunen, M., Pyšek, P., Dawson, W., Essl, F., Kreft, H., Pergl, J., Weigelt, P., Stein, A., Dullinger, S., König, C., Lenzner, B., Maurel, N., Moser, D., Seebens, H., Kartesz, J., Nishino, M., Aleksanyan, A., Ansong, M., Antonova, L.A., Barcelona, J.F., Breckle, S.W., Brundu, G., Cabezas, F.J., Cárdenas, D., Cárdenas-Toro, J., Castaño, N., Chacón, E., Chatelain, C., Conn, B., de Sá Dechoum, M., Dufour-Dror, J.-M., Ebel, A.L., Figueiredo, E., Fragman-Sapir, O., Fuentes, N., Groom, Q.J., Henderson, L., Inderjit, Jogan, N., Krestov, P., Kupriyanov, A., Masciadri, S., Meerman, J., Morozova, O., Nickrent, D., Nowak, A., Patzelt, A., Pelser, P.B., Shu, W.-s., Thomas, J., Uludag, A., Velayos, M., Verkhosina, A., Villaseñor, J.L., Weber, E., Wieringa, J.J., Yazlık, A., Zeddam, A., Zykova, E., Winter, M., 2019. The Global Naturalized Alien Flora (GloNAF) database. Ecology 100, e02542.

van Kleunen, M., Xu, X., Yang, Q., Maurel, N., Zhang, Z., Dawson, W., Essl, F., Kreft, H., Pergl, J., Pyšek, P., Weigelt, P., Moser, D., Lenzner, B., Fristoe, T.S., 2020. Economic use of plants is key to their naturalization success. Nature Communications 11, 3201.

Ward, S.F., Taylor, B.S., Dixon Hamil, K.-A., Riitters, K.H., Fei, S., 2020. Effects of terrestrial transport corridors and associated landscape context on invasion by forest plants. Biol. Invasions 22, 3051-3066.

Westoby, M., 1998. A leaf-height-seed (LHS) plant ecology strategy scheme. Plant Soil 199, 213-227.

Zhou, Q., Wang, Y., Li, X., Liu, Z., Wu, J., Musa, A., Ma, Q., Yu, H., Cui, X., Wang, L., 2020. Geographical distribution and determining factors of different invasive ranks of alien species across China. Sci. Total Environ. 722, 137929.
